# Deficiency of Hv1 Proton Channel Enhances Macrophage Antigen Presentation and Anti-Tumor Responses in Glioma

**DOI:** 10.1101/2024.10.18.619142

**Authors:** Jiaying Zheng, Lingxiao Wang, Shunyi Zhao, Katayoun Ayasoufi, Emma Goddery, Peter R. Strege, Yushan Wan, Wenjing Zhang, Manling Xie, Arthur Beyder, Aaron J. Johnson, Long-Jun Wu

**Affiliations:** Center for Neuroimmunology and Glial Biology, Institute of Molecular Medicine, University of Texas Health Science Center at Houston, Houston, Texas, USA; Mayo Clinic Graduate School of Biomedical Sciences, Rochester, Minnesota, USA; Department of Immunology, Mayo Clinic, Rochester, Minnesota, USA; Division of Gastroenterology & Hepatology, Department of Internal Medicine, Mayo Clinic, Rochester, USA; Department of Neurology, Mayo Clinic, Rochester, Minnesota, USA

**Keywords:** Voltage-gated proton channel Hv1, myeloid cells, glioma-associated macrophages, phagocytosis, antigen presentation, immune modulator, central nervous system cancer, glioblastoma

## Abstract

In the tumor microenvironment of glioblastoma, myeloid cells act as a double-edged sword—they are a major cellular component modulating the immune response while presenting a potential therapeutic target. Our study highlights the voltage-gated proton channel Hv1, predominantly expressed in myeloid cells, as a crucial regulator of their physiological functions. We discovered a strong correlation between increased Hv1 expression and poor prognosis in glioblastoma patients. Depleting Hv1 significantly extended survival in a mouse model of glioma. Employing multiple novel transgenic mouse lines, we demonstrated that Hv1 is upregulated in response to tumor presence, with glioma-associated macrophages as the principal contributors. Specifically, we identified that Hv1 in infiltrating macrophages as the major driver of survival phenotype differences. Through a combination of *in vivo* two-photon imaging, spectral flow cytometry, and spatial transcriptomic sequencing, we found that Hv1 depletion leads to reduced macrophage infiltration and enhanced antigen presentation, ultimately fostering a stronger adaptive immune response. These findings establish the Hv1 channel as a crucial new immune regulator within the brain tumor milieu, offering a promising target for reprogramming macrophage function to combat glioblastoma.

## INTRODUCTION

Protons are crucial for the physiological responses of immune cells. Central to this process is the voltage-gated proton channel Hv1, which is predominantly expressed in microglia, resident immune cells of the central nervous system (CNS), as well as in macrophages, neutrophils, B cells, and also reported in eosinophils, plasmacytoid dendritic cells, and T cell subsets ^1^. Hv1 is uniquely structured, consisting of two voltage sensor domains without a traditional pore domain ^2, 3, 4, 5, 6^. Its activation, triggered by both membrane depolarization and intracellular acidosis, typically occurs under severe physiological challenges frequently observed in pathological conditions ^7, 8^. In the periphery, Hv1 is also known to facilitating acid extrusion to trigger capacitation in human sperm ^9^, and exacerbating several cancers ^10, 11, 12^.

Hv1 has been explored for its role in supporting the production of reactive oxygen species (ROS) via NADPH oxidase 2 (NOX2) ^5, 13^. A particularly noteworthy role of Hv1 emerges following traumatic CNS injuries ^13, 14, 15, 16, 17, 18, 19, 20^; Hv1 depletion significantly alters the polarization phenotype of microglia and macrophages. These changes include reductions in ROS and cytokine production, which lead to decreased secondary tissue damage, reduced extracellular acidosis-induced neuroinflammation, and regulation of myeloid cell accumulation at CNS injury sites, highlighting Hv1 as a potent immune regulator within the CNS.

Despite these insights, the role of Hv1 in the complex immune microenvironment of tumors, especially in glioblastoma (GBM), remains largely unexplored. In GBM, myeloid cells including microglia and glioma-associate macrophages, are the unequivocally dominant immune populations ^21^ that play a dual role in the regulation of the immune response ^22, 23, 24^. Given Hv1’s critical influence on CNS injuries and our discovery of its strong correlation with poor prognosis in glioblastoma patients, our research investigates whether and how Hv1 proton channel regulate myeloid cell responses in GBM. Recognizing the limitations of existing research tools, we have developed multiple transgenic mouse lines tailored specifically for studying Hv1. Our pioneering work demonstrates the potential of targeting Hv1 to reprogram macrophages towards an anti-tumor phenotype, offering a novel and promising avenue for therapeutic intervention in glioblastoma.

## RESULTS

### Higher expression of *HVCN1* in glioma corelates with poor prognosis

To assess the significance of our studies on the Hv1 proton channel in glioma, we first analyzed its expression in samples from multiple glioma patients ^25^. The dataset included 39 contrast-enhancing glioma core samples, 36 non-enhancing FLAIR glioma margin samples, and 17 non-neoplastic brain tissue samples. We observed a significant increase in *HVCN1* expression in both tumor core and margins compared to normal brain tissues (Figure 1A). To further explore the correlation between *HVCN1* expression levels and glioma survival, we examined the TCGA dataset with a larger cohort consisting of more than 300 patients. This analysis revealed higher *HVCN1* expression in grade IV gliomas compared to grade II gliomas (Figure 1B). Consistently, by comparing the upper quartile to the lower quartile of *HVCN1* expression, we found that elevated *HVCN1* expression was associated with poorer overall survival in glioblastoma patients, indicating a correlation between *HVCN1* expression and glioma malignancy (Figure 1C).

**Figure 1.**
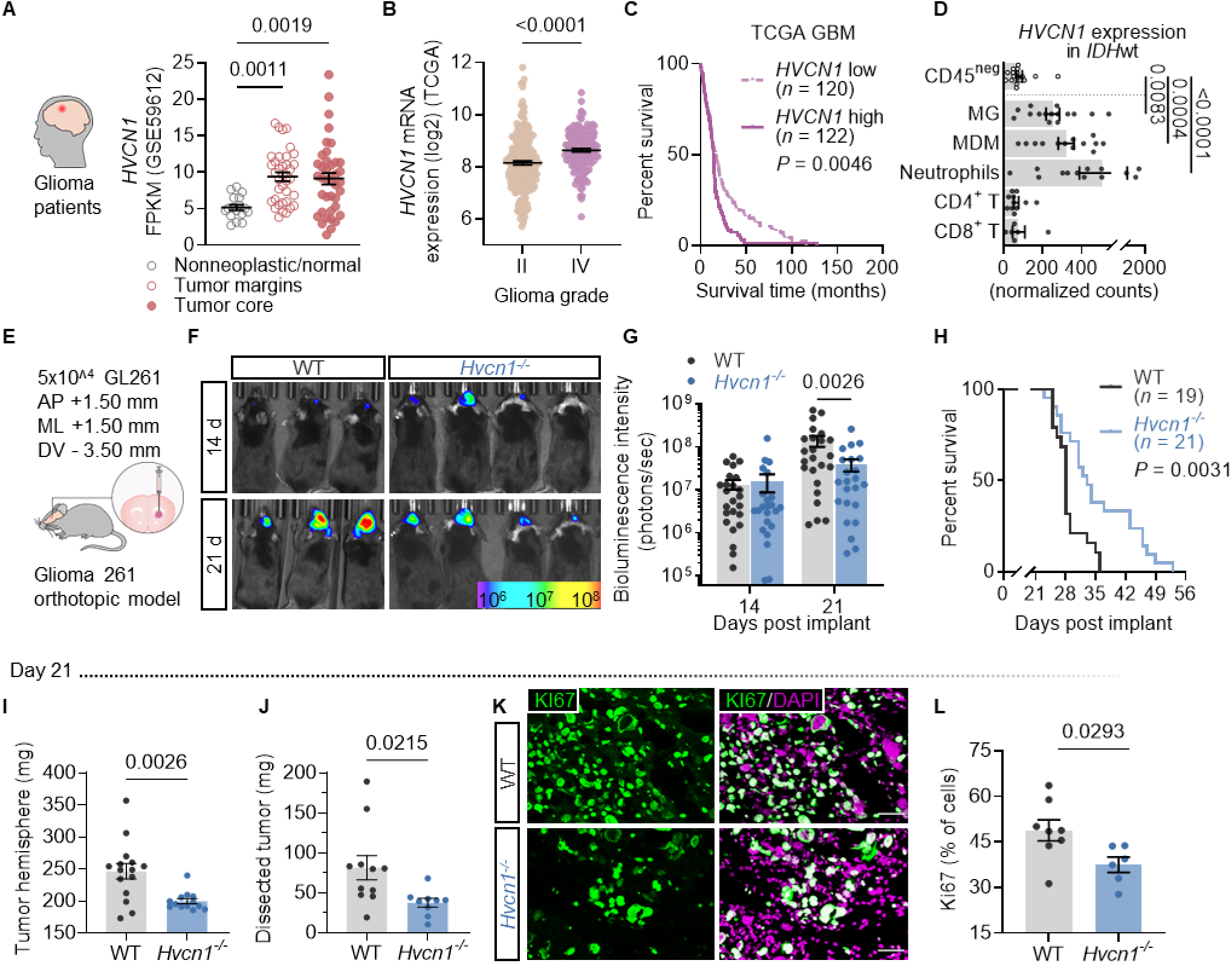
Hv1 mediates brain tumor progression in both patients and mice. A. Analysis of dataset GSE59612 revealed elevated *HVCN1* expression in both tumor margins and core compared to non-tumor tissues. B. TCGA dataset demonstrated higher *HVCN1* expression in grade IV gliomas compared to grade II. C. Survival analysis using TCGA data, with a threshold set at the 75th percentile, showed a correlation between increased *HVCN1* levels and shorter survival time in glioblastoma patients. D. In *IDH*-wt patient samples, *HVCN1* expression was predominantly observed in myeloid cells, including microglia, monocyte-derived macrophages, and neutrophils. E. Schematic representation of the mouse model. F & G. Representative and analysis of bioluminescence imaging for real-time monitoring tumor progression. At day 14, there was little difference observed between WT and *Hvcn1^-/-^*mice. However, by day 21, the tumor in *Hvcn1^-/-^* mice appeared smaller. H. Survival analysis involving 19 WT and 21 *Hvcn1^-/-^* mice demonstrates the mitigating effect of Hv1 deficiency on tumor progression. I & J. *Hvcn1^-/-^*mice exhibit smaller tumor sizes compared to WT mice 21 days post-tumor inoculation, evidenced by reduced weight of tumor hemisphere and dissected tumor. K & L. Representative images and analysis of immunostaining of KI67, a marker for proliferating cells, in day 21 tumors. A higher percentage of KI67^+^ cells was observed in WT mice. Data are presented as mean ± SEM and assessed for normal distribution using Shapiro-Wilk test. P-values were determined using two-tailed Student t-tests or two-way ANOVA for normally distributed data, and Mann-Whitney test for non-normally distributed data. Survival curves were analyzed using a log-rank test.

Next, we investigated which cell types could be the major sources of *HVCN1* in glioma. Analysis of clinical samples ^26^ revealed that myeloid cells, including microglia, monocyte-derived macrophages, and neutrophils, have the highest expression of *HVCN1* (Figure 1D). In contrast, CD45 negative cells isolated from dissected human *IDH-wt* gliomas, which likely represent the actual tumor cells, exhibit minimal *HVCN1* expression. This indicates that malignant cells are not the primary source of *HVCN1* expression; rather, elevated *HVCN1* expression in glioblastoma likely results from immune cell infiltration. This also aligns with our understanding that Hv1 proton channel is primarily expressed in the immune system.

### Hv1 deficiency slows glioma progression in the mouse model

To assess the potential impact of the Hv1 proton channel on immune responses to brain tumors, we utilized the GL261 murine brain tumor model, which is known for its extensive immune cell infiltration and aggressiveness similar to glioblastoma. We inoculated 50,000 GL261 cells into the striatum of both WT and *Hvcn1^-/-^* mice (Figure 1E). Bioluminescence imaging was used to monitor tumor growth weekly from day 14, when the tumor was established (Figure 1F). At day 14, there was no significant difference in tumor size between the two genotypes; however, by day 21, tumors in the *Hvcn1^-/-^* mice were significantly smaller (Figure 1G). Most notably, there was a survival difference between WT and *Hvcn1^-/-^* mice, with *Hvcn1^-/-^* mice showing improved survival (Figure 1H).

Further pathological analysis of brains taken 21 days post-inoculation yielded consistent results that *Hvcn1^-/-^* mice exhibit slower tumor growth compared to WT counterparts. Since GL261 tumor cells do not cross hemispheres, weighing the tumor-burdened hemisphere provides an estimate of tumor size. In line with the bioluminescence results, the *Hvcn1^-/-^* mice had lighter tumor-burdened hemispheres compared to the WT group (Figure 1I). Weighing dissected tumors showed a similar pattern, with tumors from *Hvcn1^-/-^* mice being lighter (Figure 1J). In addition, proliferation staining using KI67 revealed a higher percentage of proliferating cells in the tumor regions of WT mice compared to *Hvcn1^-/-^* mice (Figure 1K &1L), indicating that the WT environment is more favorable for faster tumor growth. Taken together, our results suggest that Hv1 deficiency slows tumor progression in the mouse model of glioma.

### Profile Hv1 expression during glioma progression using a novel transgenic mouse model

We then investigated the Hv1 expression pattern during glioma progression. Previous literature establishes that the Hv1 channel primarily resides within multiple immune cell populations ^1^. However, determining which cell types express Hv1 and how its expression is regulated in response to tumors presents two significant challenges: the lack of high-quality commercially available antibodies for Hv1 detection and the very low RNA expression levels of *Hvcn1* at the single-cell level. Most previous validations of Hv1 rely on electrophysiology, which has limitations in *in vivo* diseased models and makes it difficult to identify the multiple potential immune cell type candidates.

To this end, we engineered a novel transgenic mouse model (Figure 2A) in which the Hv1 protein is directly tagged with a hemagglutinin (HA) tag. This tag enables precise localization of Hv1 through HA staining (Figure 2B) and may facilitate Hv1 pull-down assays for further research. Additionally, we introduced an mCherry fluorophore downstream of the Hv1 promoter, separated by a P2A linker, allowing for simultaneous visualization of Hv1 expression. Validation in HEK 293 cells demonstrated that after transfection with the plasmid *Hvcn1-HA-P2A-mCherry*, evidenced by both mCherry and HA expression (Figure 2C & 2D), the cells exhibited functional voltage-gated proton signature currents, whereas non-transfected cells displayed no currents (Figure 2E - 2F’).

**Figure 2.**
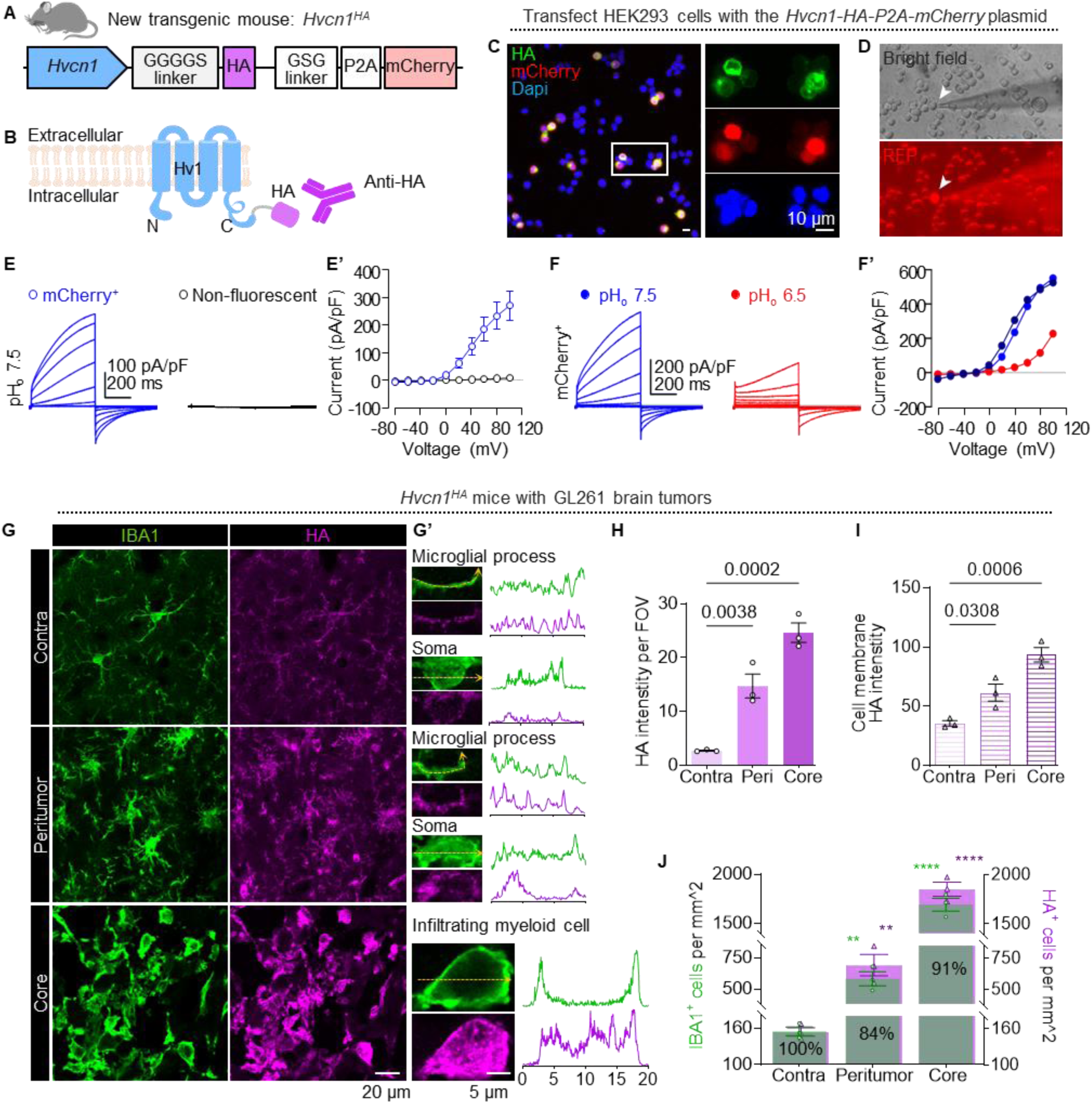
Characterize Hv1 expression profile during glioma progression using a novel transgenic mouse model. A. Diagram of the novel transgenic mouse model. B. The diagram displays the location of the HA tag and the concept behind using anti-HA antibodies for Hv1 protein detection or future pull-down assays. C. Immunostaining revealed that HA and mCherry signals were colocalized in the same cells, confirming successful transfection. D. Cells transfected with the plasmid displayed mCherry^+^ signals during patch-clamp recording. E - F’. Only cells exhibiting mCherry^+^ signals have Hv1 signature current, indicating its specific expression in HEK293 cells. The Hv1 proton channel in transfected cells was functional and exhibited the reported characteristics, validating the effectiveness of the plasmid. In F’, *n* = 11 non-fluorescent cells at pHo = 7.5, *n* = 7 rinses with pHo = 6.5, *n* = 3 washouts to pHo = 7.5. G. Representative image illustrating the staining of myeloid cells with IBA1 and HA. Hv1 proton channel expression is visualized through HA tag staining in the contralateral hemisphere, peritumoral region, and the tumor core (scale bar: 20 μm). G’. Inset of G showing processes and somas of IBA1^+^ cells and their Hv1 expression. The line graph depicts the intensity of IBA1 (green) and HA tag (magenta) under the same confocal settings (scale bar: 5 μm). H. Overall HA intensity per field of view showing Hv1 is more abundant in tumor core and peritumoral region than contralateral region. I. HA intensity per cell, showing the cells elevated Hv1 expression in the tumor core and peritumoral regions compared to the contralateral hemisphere. J. Colocalization analysis between HA and IBA1 demonstrates that most HA signals overlap with IBA1^+^ cells. Data are presented as mean ± SEM. P-values were determined using one-way ANOVA.

In the naïve *Hvcn1-HA-mCherry* mouse brains, HA is specifically colocalized with IBA1^+^ cells, consistent with previous research indicating that Hv1 is predominantly expressed by microglia in the CNS ^7, 13, 14, 18^ (Supplementary figure 1A). The fine HA staining observed on the soma membrane and processes clearly indicates the abundance of Hv1 protein on the plasma membrane. Surprisingly, we did not detect any visible mCherry signals in microglia. This discrepancy could be due to low *Hvcn1* RNA expression, which is consistently observed in the single-cell dataset, and results in the promoter not being robust enough to drive mCherry expression in microglia. However, the underlying reasons for this low gene expression remain unexplored.

When the tumor is present, the HA staining intensity is stronger in the tumor regions than in the contralateral hemisphere, suggesting an upregulation of Hv1 expression in response to the tumor (Figure 2H and Supplementary Figure 1C). However, the mCherry signal remained undetectable in the contralateral hemisphere and was faint in IBA1^+^ cells in both the peritumoral and tumor core regions (Supplementary Figure 1B). Nevertheless, detailed cellular analysis revealed that HA staining in the processes and soma of peritumoral microglia is significantly enhanced compared to microglia in the contralateral hemisphere. In the core regions, myeloid cells exhibit saturated HA staining intensity under the same confocal settings (Figure 2G, 2G’, and 2I). This staining is not confined to the membrane surface but extends into the cytosol, indicating active translation from mRNA or recruitment of the channel to other cellular organelles, such as early endosomes, upon cellular activation ^27^. Further localization analysis of HA and IBA1 revealed that HA expression predominantly overlaps with IBA1^+^ cells (Figure 1J). Taken together, using this novel mouse model, we demonstrate that Hv1 is upregulated in response to the tumor, with microglia and glioma-associated macrophages identified as the primary source of Hv1.

### Hv1 promotes myeloid cell infiltration in glioma

Given that the upregulated expression of Hv1 proton channel in tumor hemispheres is primarily in myeloid cells, we next investigated whether Hv1 depletion alters myeloid cell dynamics. Tumor hemispheres were collected on day 21 after tumor inoculation for flow cytometry analysis (Figure 3A). The t-distributed stochastic neighbor embedding (t-SNE) map displayed major immune cell clusters (CD45^+^) identified through flow cytometry, with infiltrating myeloid cells representing the largest population, followed by microglia, CD4^+^ T cells, NK cells, CD8^+^ T cells, and B cells (Figure 3B). Excitingly, the largest population, infiltrating myeloid cells, identified as CD11b^high^, CD45^high^, TCRβ^negative^, B220^negative^, and NK1.1^negative^, showed a significant difference between WT and *Hvcn1^-/-^* mice. In WT mice, infiltrating myeloid cells accounted for approximately 70% of all immune cells at this time point. Remarkably, Hv1 deficiency reduced this population to 46% (Figure 4B’). Statistical analyses of both the percentage of infiltrating myeloid cells among all immune cells and the actual cell counts confirmed the decrease in infiltrating myeloid cells in *Hvcn1^-/-^*mice (Figure 3C & 3D). Importantly, these infiltrating myeloid cells consist mostly of monocytes and macrophages ^21^.

**Figure 3.**
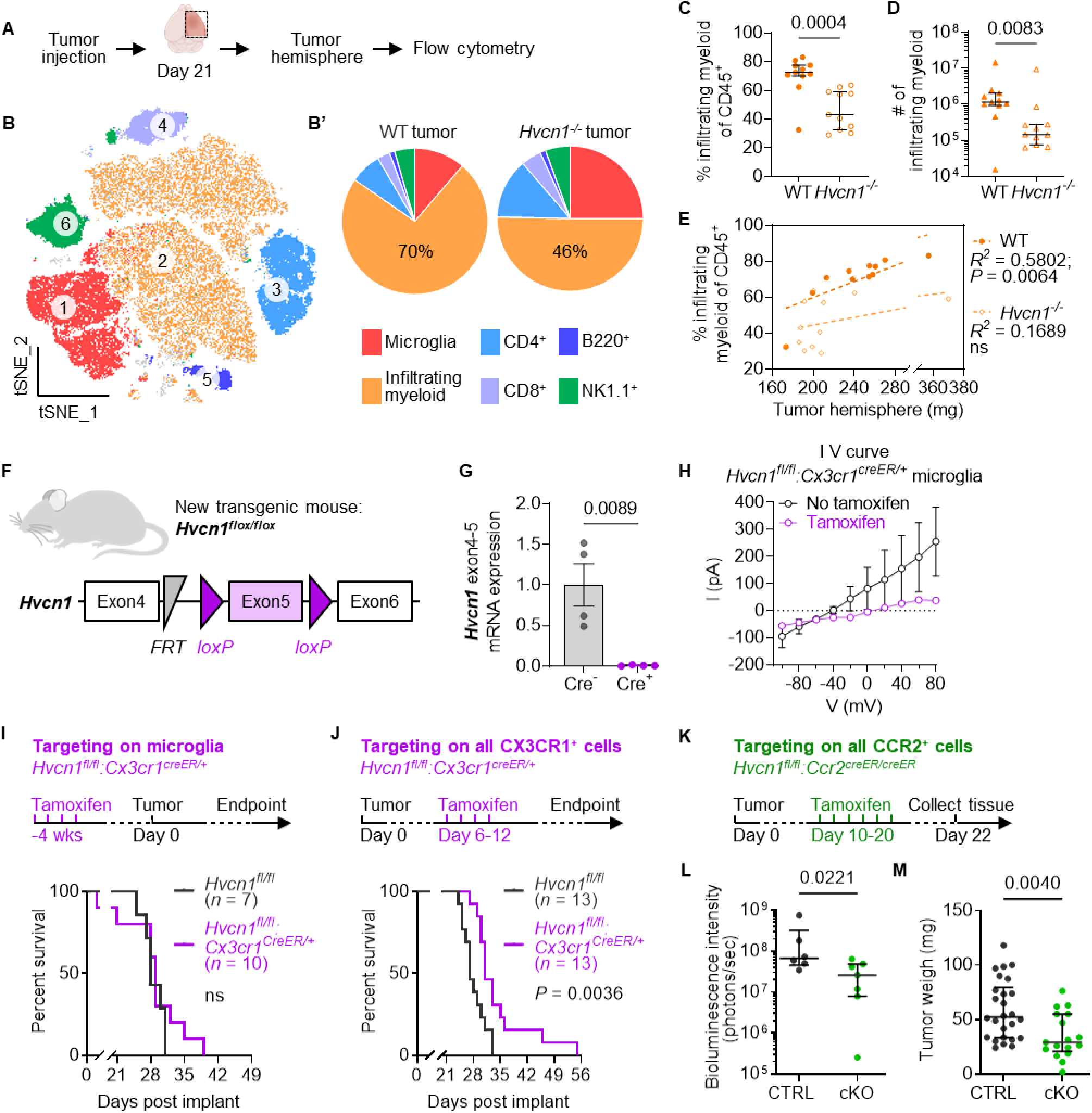
Hv1 depletion impacts mostly infiltrating myeloid cells, and this population is the major contributor drives the tumor progression difference between genotypes. A. Experiment design diagram. B. tSNE map illustrating all major immune cell clusters detected by spectrum flow cytometry in the tumor hemispheres. B’. A pie chart shows the dominant immune population in WT glioma is infiltrating myeloid cells, which are reduced in *Hvcn1^-/-^* mice. C & D. Analysis of the percentage of infiltrating myeloid cells among CD45^+^ cells and the actual cell count of infiltrating myeloid cells showing the reduction of infiltrating myeloid cells in *Hvcn1^-/-^* mice. E. Correlation analysis reveals a strong association between tumor burden and the percentage of infiltrating myeloid cells in WT, indicated by a high R² value and significant P value. In contrast, *Hvcn1^-/-^* mice show a minimal correlation with a low R² value and nonsignificant P value. F. Diagram of new transgenic mouse line. G. Validation of the Hv1 knockout efficiency via qRT-PCR on sorted microglia from naïve *Hvcn1^flox/flox^* and *Hvcn1^flox/flox^:Cx3cr1^creER/+^*mice treated with tamoxifen. H. Functional validation through electrophysiology on brain slices confirms the absence of Hv1 currents in *Hvcn1^flox/flox^:Cx3cr1^creER/+^*microglia after tamoxifen administration. I. Investigation of microglial Hv1 proton channel depletion in the tumor model shows no survival difference between *Hvcn1^flox/flox^:Cx3cr1^creER/+^* mice and their littermates *Hvcn1^flox/flox^*. J. Examining the impact of depleting the Hv1 proton channel in all CX3CR1^+^ cells during early tumor progression; a survival benefit is observed in the conditional knockout group. K. Experimental design focuses on depleting the Hv1 proton channel primarily in infiltrating myeloid cells by crossing *Hvcn1^flox/flox^*with *Ccr2^creER-Gfp^*. L. *In vivo* bioluminescence imaging reveals that Hv1 depletion in infiltrating myeloid cells leads to smaller tumors. M. Dissected tumor tissues also show reductions in the conditional knockout group. Data were assessed for normal distribution using Shapiro-Wilk test are presented as mean ± SEM for normally distributed data or medium ± interquartile range for non-normally distributed data. P-values were determined using two-tailed Student t-tests for normally distributed data, and Mann-Whitney test for non-normally distributed data. Simple linear regression with Pearson correlation coefficients was used for correlation analysis. Survival curves were analyzed using a log-rank test.

**Figure 4.**
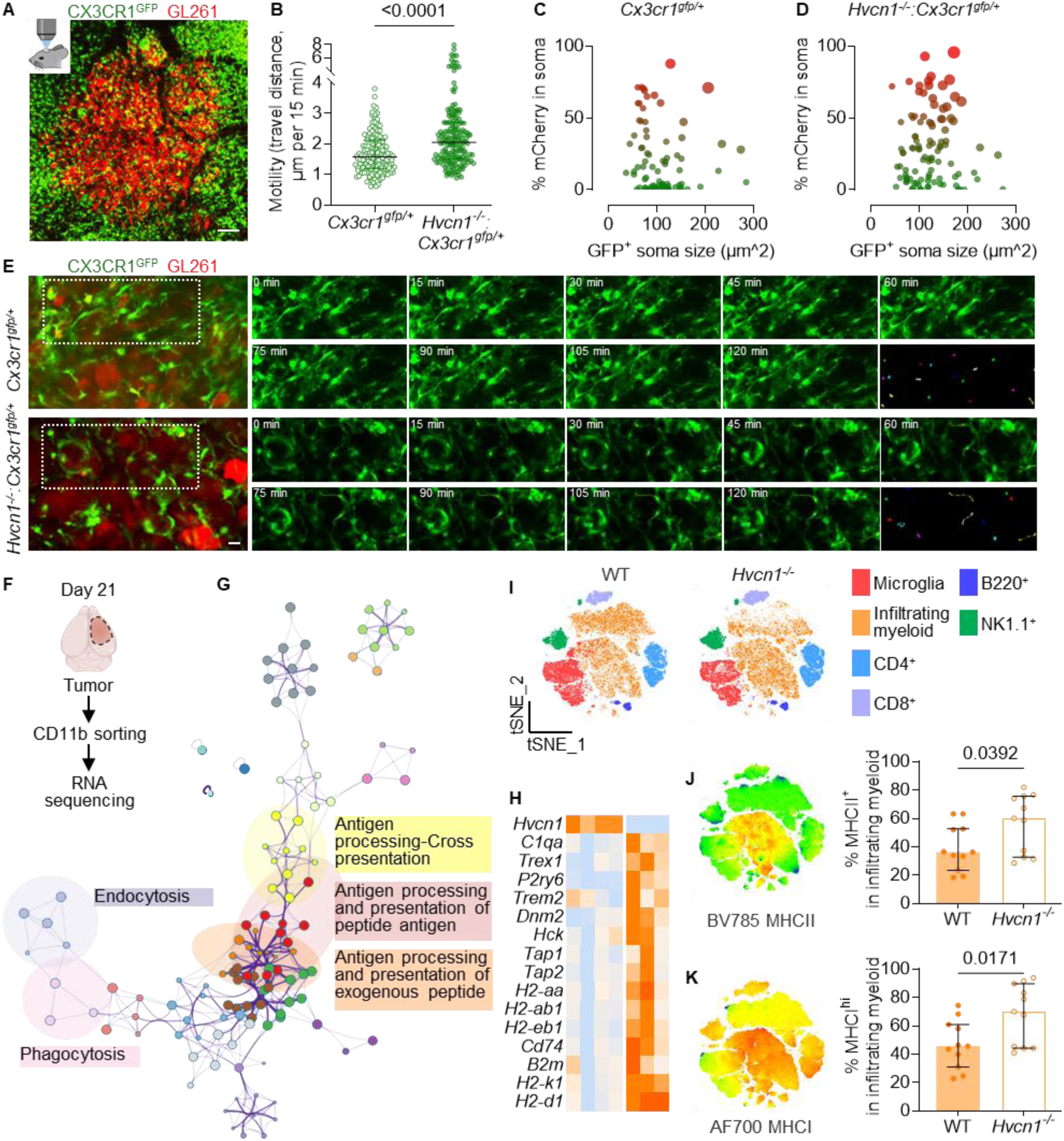
Hv1 depletion enhances tumor cell phagocytosis in infiltrating myeloid cells, leading to improved antigen presentation. A. Representative two-photon tiling imaging captures interactions between tumor cells (red) and CX3CR1+ cells (green). B. The moving speed of CX3CR1^+^ cell somas is calculated by measuring the average distance moved every 15 minutes over a 2–3-hour period. C & D. Analysis of myeloid cell size and mCherry content shows variations in mCherry signal intensity across different sizes of myeloid cells. Notably, *Hvcn1^-/-^*:*Cx3cr1^gfp/+^*cells display increased mCherry signals, suggesting higher uptake of tumor debris. E. Representative images of tracing the movement of CX3CR1^+^ cells within the tumor. F. Diagram of the procedure for dissecting the tumor and sorting CD11b^+^ cells for RNA sequencing. G A functional clustering network that illustrates the relationships between the top 20 enriched terms generated based on DEGs in *Hvcn1^-/-^*. Pathways with a similarity score above 0.3 are connected by lines and color-coded, with key pathways highlighted and labeled. H. Heatmap showing expression levels of selected genes from highlighted clusters in the GO pathway analysis, with orange indicating higher expression levels. I. tSNE map illustrating major immune cell populations in both wild-type and *Hvcn1^-/-^* tumor hemispheres. J & K. tSNE map displaying the expression of MHC class II and MHC class I. Analysis reveals that percentage of MHC class II and MHC class I among myeloid cells both increase in *Hvcn1^-/-^*. Data were assessed for normal distribution using Shapiro-Wilk test are presented as mean ± SEM for normally distributed data or medium ± interquartile range for non-normally distributed data. P-values were determined using two-tailed Student t-tests for normally distributed data, and Mann-Whitney test for non-normally distributed data.

To ensure that the differences in infiltration were not due to variations in tumor size, we plotted the weight of tumor hemispheres against the percentage of infiltrating myeloid cells among CD45^+^ cells. In WT mice, a strong positive correlation was observed, indicating that higher tumor burden led to a greater percentage of infiltrating myeloid cells. However, this correlation was absent in *Hvcn1^-/-^* mice (Figure 3E), further highlighting the impact of Hv1 deficiency on this cell population.

### Hv1 in infiltrating glioma-associated macrophages contribute to tumor progression

To further investigate whether the Hv1 proton channel in myeloid cells contributes to differences in tumor progression, we generated a transgenic mouse that enables a conditional knockout of Hv1 in our cells of interest. A loxP site was inserted into exon 5, allowing for gene deletion in the presence of CRE recombinase (Figure 3F). The knockout efficiency was confirmed by qRT-PCR using sorted microglia from naïve mouse brains of *Hvcn1^flox/flox^* and *Hvcn1^flox/flox^:Cx3cr1^creER/+^*mice treated with tamoxifen (Figure 3G). Additionally, the knockout efficiency was validated functionally through electrophysiology, which confirmed the absence of Hv1 currents in *Hvcn1^flox/flox^:Cx3cr1^creER/+^* microglia after tamoxifen administration (Figure 3H).

We first targeted microglial Hv1. Tamoxifen was administered 4 weeks prior to tumor inoculation to induce exon 5 deletion of *Hvcn1* in CX3CR1^+^ cells. Since microglia are long-lived and self-replenishing, after 4 weeks, they will continue to carry the *Hvcn1* exon 5 deletion, while most peripheral immune cells will have undergone turnover and been replenished with unmodified genes. Interestingly, we found no survival difference between *Hvcn1^flox/flox^:Cx3cr1^creER/+^* mice and their littermates *Hvcn1^flox/flox^* (Figure 3I). This indicates that depletion of Hv1 specifically in microglia may not be sufficient to delay robust tumor progression.

Next, we targeted all CX3CR1^+^ cells during tumor progression to include both microglia and a large proportion of infiltrating monocytes/macrophages ^28, 29^. Tamoxifen was administered from day 6 to day 12 after tumor inoculation, corresponding to the early phase of tumor initiation and myeloid cell response. This strategy demonstrated a survival benefit in the conditional knockout group of *Hvcn1^flox/flox^:Cx3cr1^creER/+^* mice, suggesting that the survival difference requires infiltrating CX3CR1^+^ cells, primarily glioma-associated macrophages (Figure 3J).

To further confirm the importance of Hv1 in glioma-associated macrophages, we crossed *Hvcn1^flox/flox^* with *Ccr2^creER-Gfp^* mice, as most infiltrating monocytes/macrophages from the periphery are CCR2 signaling dependent ^30^. Tamoxifen was administered from day 10 to day 20, and tissues were collected on day 22 (Figure 3K). By day 22, *in vivo* bioluminescence imaging showed that the conditional knockout group exhibited dimer signals, indicating smaller tumors (Figure 3L). Consistently, dissected tumor weights were also reduced in the conditional knockout group (Figure 3M). Together, these results demonstrate that Hv1 in glioma-associated macrophages contributes to tumor progression.

### *In vivo* two-photon imaging reveals that Hv1 depletion increases the motility of glioma-associated macrophages

We next investigated how Hv1 influences the behavior of glioma-associated macrophages. MCherry-GL261 tumor cells ^31^ were inoculated into the somatosensory cortex at a density of approximately 3,000 cells to prevent overly aggressive growth and allow for extended observation under the imaging window. After a week, we observed the establishment of the tumor using *in vivo* two-photon tiling imaging, with mCherry tumor cells (red) forming a dense cellular cluster in the central area of the window, and a significant number of GFP^+^ myeloid cells (green) actively responding to and infiltrating the tumor (Figure 4A).

In the tumor core, we observed two main types of GFP^+^ cells: one type appeared small and round, rolling in a tunnel-like shape (possibly associated with vasculature) and moving rapidly, which could represent monocytes or CX3CR1^+^ T cells. The other type displayed an ameboid shape with a few pseudopodia, moving more slowly and resembling glioma-associated macrophages. To observe these slower-moving macrophages, we acquired images every 15 minutes over a few hours and traced their cell somas for motility analysis. We found that *Hvcn1^-/-^*:*Cx3cr1^gfp/+^* cells exhibited higher motility, with more cells actively crawling (Figure 4B & 4E).

It is worth noting that when analyzing the motility of myeloid cells in the tumor environment, their speed may be influenced by vascular pulsation and tumor cells. Therefore, even though some cells of interest remained stationary for most of the time during the 2-3-hour timeframe, the speed analysis typically will not yield a value of zero. For macrophages moving with directed purpose, we observed their pseudopodia extending rapidly, anchoring to the surrounding environment, and following this initial extension, the cell pulls its cell soma forward toward the leading edge. This behavior was more frequently observed in the *Hvcn1^-/-^*:*Cx3cr1^gfp/+^*cells than that from control *Cx3cr1^gfp/+^* cells, contributing to the higher motility after Hv1 deficiency.

Additionally, we noted multiple macrophages merging pseudopodia to form a ring-like structure around tumor cells, actively encircling the tumor and resulting in the rupture of those tumor cells (Figure 4E). This type of structure was more frequently observed in the *Hvcn1^-/-^*:*Cx3cr1^gfp/+^* group. However, due to the limited timeframe, tracing its formation and consequences proved challenging, leaving the detailed mechanism unknown.

In addition to the rupture of tumor cells, we also noted glioma-associated macrophages taking up mCherry^+^ debris in their soma, consistent with our previous findings ^31^. Further analysis showed that *Hvcn1^-/-^*:*Cx3cr1^gfp/+^* cells contained a higher percentage of mCherry^+^ debris within their somas compared to the *Cx3cr1^gfp/+^* cells (Figure 4C & 4D). This observation leads us to consider one of the increasingly recognized anti-tumor functions of macrophages: phagocytosis and antigen presentation.

### Hv1 depletion enhances the phagocytosis and antigen presentation of glioma-associated macrophages

We further explored how Hv1 modulates the transcriptomic profile of glioma-associated macrophages using RNA sequencing. In well-established tumors, microglia and glioma-associated macrophages occupy distinct locations ^32, 33, 34^; microglia form a barrier surrounding the tumor, while infiltrating monocytes and macrophages are primarily localized within the tumor itself. Based on this observation, we dissected the tumor from the tumor hemispheres on day 21 and collected CD11b^+^ cells using magnetic-activated cell sorting. These cells are theoretically composed primarily of infiltrating myeloid cells (Figure 4F). Selected marker genes confirmed that the sorted cell populations were predominantly infiltrating myeloid cells, characterized by high expression of *Cx3cr1*, *Aif1* (encoding IBA1), *Itgam* (encoding CD11b), *Trem2*, *Cd14*, *Lyz2*, and *Cd68*, while exhibiting low expression of microglial markers such as *Tmem119* and *P2ry12*.Additionally, these cells demonstrated low or absent RNA expression of B cell marker genes (*Cd19*, *Mzb1*, *Ms4a1*), T cell markers (*Cd3d*, *Cd8a*, *Gzmk*, *Cd4*), markers for NK cells and subsets of T cells (*Cd2*, *Cd160*, *Ncam1*, *Klrb1c* encoding NK1.1), as well as low expression of dendritic cell markers (*Batf3*, *Flt3*, *Clec9a*, *Xcr1*). They also showed no expression of mast cell and basophil markers (*Ms4a2*) and neutrophil markers (*Ly6G*) (Supplementary Figure 2A).

We first performed Gene Set Enrichment Analysis (GSEA) to identify biologically relevant gene sets that exhibit significant changes upon depletion of the Hv1 proton channel. This approach enables us to capture coordinated changes in pathways, minimizing the noise associated with individual differentially expressed genes (DEGs). Our findings indicate that in *Hvcn1^-/-^* mice, GSEA pathways reveal a strong myeloid activation involving antigen presentation and regulating immune signaling. This suggests that these myeloid cells are more primed to recognize and respond to antigens, potentially facilitating a more effective adaptive immune response against tumors (Supplementary Figure 2B).

Building on the significant pathways identified through GSEA, we proceeded to analyze the DEGs for a more focused exploration using Gene Ontology (GO) analysis. By setting a threshold of FPKM > 50 and adjusting the p-value to < 0.05, we identified a total of 465 DEGs, with 57 genes upregulated in the wild-type (WT) group and 408 genes upregulated in the *Hvcn1^-/-^*mice. We then input the 408 upregulated DEGs into Metascape for GO pathway analysis, generating a functional clustering network that illustrates the relationships between the enriched terms. A subset of these enriched terms was selected and represented as a network plot, with terms having a similarity score greater than 0.3 connected by lines and color-coded for clarity. Consistent with our GSEA analysis, the top-enriched clusters identified were associated with macrophage anti-tumor responses, encompassing key processes such as endocytosis, phagocytosis, and antigen processing and presentation (Figure 4G). The corresponding genes for these highlighted clusters were selected and listed (Figure 4H).

The enhanced antigen expression was further validated at the protein level through flow cytometry. Among the six major immune populations identified, the cluster of infiltrating myeloid cells (mostly glioma-associated macrophages) exhibited a reduction in the tSNE map for the *Hvcn1^-/-^* group (Figure 4I). Notably, at this time point, infiltrating myeloid cells in the *Hvcn1^-/-^* mice showed a decreased proportion of F4/80^high^ Ly6C^low^ SSC^high^ cells, while there was an increase in cells characterized by a Ly6C^high^ F4/80^low^ SSC^low^ phenotype (Supplementary Figure 2C & 2D). Interestingly, MHC class II expression was primarily observed in the subset defined by elevated Ly6C and moderate levels of F4/80, meanwhile MHC class I expression was also higher in this same subset of myeloid cells (Figure 4J & 4K). Consequently, quantification of MHC class II and MHC class I expression revealed that *Hvcn1^-/-^* mice had a higher percentage of infiltrating myeloid cells expressing both MHC class II and MHC class I compared to WT (Figure 4J & 4K). Together, these findings demonstrate that Hv1 deficiency significantly modulates glioma-associated macrophages towards a more anti-tumor response, particularly enhancing their antigen presentation capabilities.

### Spatial transcriptomic analysis reveals enhanced adaptive immune response after Hv1 deletion

To gain deeper insights into the spatial response differences between WT and *Hvcn1^-/-^* mice, we employed spatial RNA sequencing (10x Visium) to create transcriptomic maps across tumor and peritumoral brain regions. Coronal sections from both WT and *Hvcn1^-/-^*mice were collected three weeks post-tumor inoculation, aligning with previous time points. The transcriptomic profiles were then clustered and visualized using UMAP, with each cluster mapped to anatomical brain regions defined by the Allen Mouse Brain Atlas ^35^ (Figure 5A & 5B). These clusters displayed regional segregation of brain tissues and were consistent with histological annotations, including the anterior cortical amygdaloid nucleus (ACo), cerebral cortex (CTX), ependymal layer, leptomeninges, pallidum (PAL), and striatum (STR). Most importantly, tumor-related regions, including the tumor core and peritumoral region, were distinctly clustered away from other brain areas and closely matched the dense DAPI staining that marks the tumor (supplementary figure 3A). All clusters were present in both genotypes; however, the *Hvcn1^-/-^*tumors were smaller, resulting in smaller peritumoral and tumor core clusters being represented on the UMAP.

**Figure 5.**
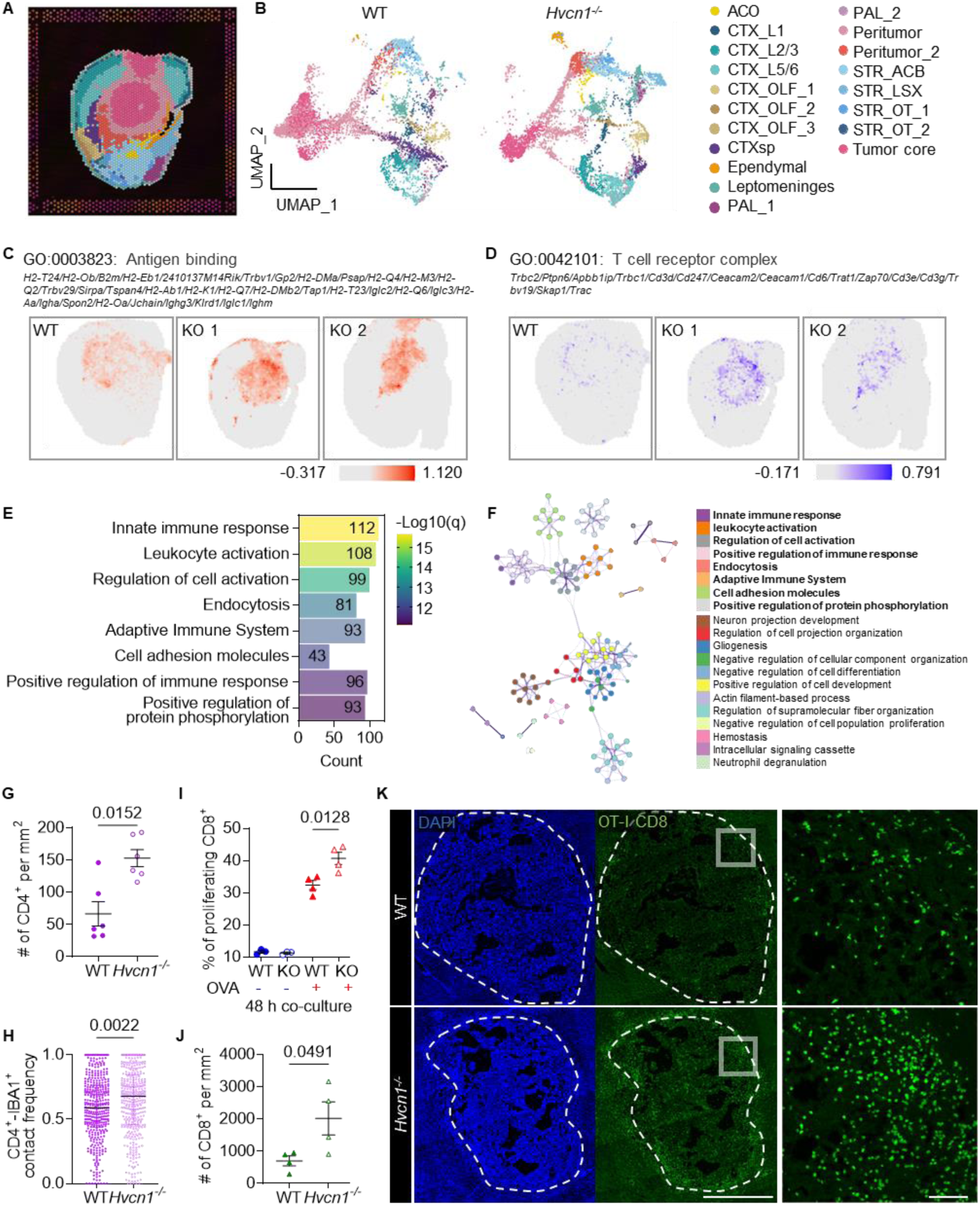
Enhanced adaptive immune response following innate immune activation in *Hvcn1^-/-^* mice. A & B. The transcriptomic profiles from spatial RNA sequencing are clustered and visualized using UMAP. Each cluster is mapped to anatomical brain regions as defined by the Allen Mouse Brain Atlas. C & D. Color gradient plots show the scores of antigen binding and T cell receptor complex across tissue sections. E. Selected top 20 GO pathways based on DEGs in *Hvcn1^-/-^* mice. F. A functional clustering network that illustrates the relationships between the top 20 enriched terms generated based on DEGs in *Hvcn1^-/-^*. G. Quantification of CD4^+^ T cells per area of interest in the tumor, showing its increased infiltration in *Hvcn1^-/-^*. H. Analysis based on over 60 cells per animal across a total of six animals per group shows elevated contact frequency between CD4^+^ T cells and IBA1^+^ myeloid cells in the tumor regions of *Hvcn1^-/-^*. I. *In vitro* antigen presentation experiment demonstrates that *Hvcn1^-/-^* BMDM can more efficiently process ovalbumin (OVA) and present it to OT-1^+^ CD8^+^ T cells. J. Quantification of GFP^+^ OT-1^+^ CD8^+^ T cells per area in the tumor region, indicating a more antigen-dependent CD8^+^ T cell infiltrate in *Hvcn1^-/-^* (scale bar: 1000 µm; inset scale bar: 100 µm). Data were assessed for normal distribution using Shapiro-Wilk test are presented as mean ± SEM for normally distributed data or medium ± interquartile range for non-normally distributed data. P-values were determined using two-tailed Student t-tests for normally distributed data, and Mann-Whitney test for non-normally distributed data.

Our primary focus is on the tumor and the surrounding peritumoral regions. We began by GSEA on predefined gene sets to investigate the biological processes associated with the previously identified differences in antigen presentation. The GSEA results for the tumor core highlighted several immune-related pathways enriched in *Hvcn1^-/-^*, including antigen binding, T cell receptor complex, antigen processing and presentation of peptide antigens via MHC Class II, and immunoglobulin complexes (Supplementary Figure 3B). Similarly, in the peritumoral region, pathways related to antigen binding and the T cell receptor complex were also prominent in the *Hvcn1^-/-^* group (Figure 5C & 5D). Spatial analysis revealed that antigen binding predominantly filled the entire tumor area, whereas the T cell receptor complex was mainly located in the tumor area, with a higher concentration near the tumor border.

While GSEA focuses on the collective behavior of gene sets and provides a broader view of biological processes, it can be further complemented by analyzing DEGs through GO pathway analysis. Notably, both the tumor core and peritumoral region exhibited the highest number of DEGs between WT and *Hvcn1^-/-^*, with 2,826 DEGs identified in the peritumoral region and 2,708 in the tumor core (Supplementary Figure 3C). Importantly, the DEGs were not confined to these tumor-adjacent areas; all other brain regions displayed differential expression as well, with the number of DEGs ranging from 1 to 1,963.

Given the significant differences in gene expression observed in the tumor core area and considering that our previously identified main phenotypic differences occur in this region, we performed GO pathway analysis on the 1,302 DEGs that were expressed at higher levels in *Hvcn1^-/-^*compared to WT. Among the top 20 listed pathways, we identified there 112 genes involved in innate immune response, 108 genes involved in leukocyte activation, 99 genes involved in regulation of cell activation, 81 genes involved in endocytosis, 93 genes involved in adaptive immune system, 43 genes involved in cell adhesion molecules, 96 genes involved in positive regulation of immune response, and 93 genes involved in positive regulation of protein phosphorylation (Figure 5E). These 20 pathways were further illustrated in a functional clustering network (Figure 5F). Overall, the findings from both the GSEA and the GO analysis of DEGs indicate that the immune response in *Hvcn1^-/-^* mice is significantly enhanced, particularly highlighting the enhanced T cells responses to the more active antigen presentation by macrophages.

### Hv1 depletion boosts antigen-mediated responses in both CD4^+^ and CD8^+^ T cells

We next validated the enhanced adaptive immune response, with a particular focus on T cell responses. Notably, recent studies including ours have highlighted the significance of the MHC class II-CD4^+^ T cell pathway ^36^, ^3731^.To further investigate this, we examined CD4^+^ T cells in tumor regions and found increased infiltration of CD4^+^ T cells in *Hvcn1^-/-^* mice (Figure 5G). More importantly, after analyzing over 60 cells per animal across a total of six animals per group, we observed an elevated contact frequency between CD4^+^ T cells and IBA1^+^ myeloid cells in the tumor regions of *Hvcn1^-/-^*mice (Figure 5H). The increased infiltration of CD4^+^ T cells and the enhanced contact suggest that the heightened MHC class II antigen expression in *Hvcn1^-/-^*mice may functionally induce CD4^+^-mediated immunity.

While the recognition of CD4^+^ anti-tumor effects is increasing, cytolytic CD8^+^ T cells are known to play a crucial role in tumor killing primarily MHC class I dependent. Given that we observed significant increases not just MHC class II but also MHC class I expression in *Hvcn1^-/-^* myeloid cells, we next evaluated the CD8^+^ T cell response, focusing on MHC class I-mediated activation. We first performed a classic *in vitro* antigen presentation assay. Bone marrow-derived macrophages (BMDMs) were collected from both WT and *Hvcn1^-/-^*, treated with ovalbumin (OVA), and stimulated with lipopolysaccharide (LPS) before being co-cultured with OT-1^+^ CD8^+^ T cells for 48 hours. OT-1^+^ T cells are transgenic T cells that express a T cell receptor (TCR) specific for the OVA^257-264^ peptide derived from OVA, presented by MHC class I molecules. When BMDMs process OVA, they display this peptide on their MHC class I molecules, enabling recognition by OT-1^+^ CD8^+^ T cells. Upon binding to the peptide-MHC class I complex, OT-1^+^ CD8^+^ T cells become activated, resulting in T cell proliferation. Indeed, in the OVA-treated group, we observed significant proliferation of CD8^+^ T cells compared to the non-treated group, indicating that this proliferation is antigen-dependent. Notably, *Hvcn1^-/-^* BMDMs promoted greater proliferation of CD8^+^ T cells. These results suggest that Hv1 deficiency in BMDMs enhanced their cross-antigen presenting capabilities (Figure 5I).

To further confirm this phenotype in the context of glioma, we conducted an *in vivo* antigen presentation assay. We transplanted an equal number of GFP^+^ OT-1^+^ CD8^+^ T cells into either WT or *Hvcn1^-/-^* mice with GL261-OVA inoculation. By day 21, we collected the tumor hemispheres and found that GFP^+^ OT-1^+^ CD8^+^ T cells had infiltrated both the tumor core and border areas. This infiltration suggests that myeloid cells have taken up tumor cells to some extent, processed the OVA, and presented the OVA^257-264^ peptide via MHC class I for T cell recognition (Figure 5J). Furthermore, consistent with our *in vitro* data, the number of GFP^+^ OT-1^+^ CD8^+^ T cells was significantly higher in *Hvcn1^-/-^* mice compared to WT, demonstrating that cross-presentation was enhanced in the *Hvcn1^-/-^* myeloid cells (Figure 5K). Collectively, these findings underscore the role of the Hv1 proton channel in modulating key immune-related pathways in glioma-associated macrophages and the subsequent adaptive immune response in the tumor progression.

## DISCUSSION

We demonstrated that elevated expression of *HVCN1* in glioma not only correlates with poorer prognosis but also accelerates tumor progression. One of key innovations of our study is the development of novel transgenic mouse lines to enrich the toolkit in studying Hv1 proton channel in vivo. The HA-tagged Hv1 proton channel mouse line allowed us to precisely pinpoint Hv1 protein expression pattern, revealing that its levels increase in response to glioma progression. It would further provide a precise isolation of Hv1 using the HA tag, facilitating the screening of multiple antibodies that can target Hv1, and identify potential proteins have direct interactions with Hv1 in different physiology or pathological conditionals. Thus, the mouse line is extremely useful for studying Hv1 proton channel, circumventing the caveats of low Hvcn1 RNA expression and the lack of effective antibodies. In addition, the generation of Hv1-floxed mouse line allows us to interrogate the role of Hv1 in specific cells in health and disease. By integrating multiple Cre lines in the current study, we demonstrated that Hv1 in infiltrating myeloid cells is a key driver of immune suppression during tumor progression.

Hv1 deficiency, leading to reduced macrophage accumulation, may provide a survival advantage in brain tumors. This reduction aligns with previous findings in traumatic CNS injuries ^14, 38^. Excessive infiltration and accumulation of these cells is typically associated with immunosuppression, as they can inhibit T cell functions through mechanisms such as the expression of inhibitory checkpoints and the release of suppressive cytokines ^39^. Excessive ROS can induce immunosuppression ^40^, and Hv1 plays a crucial role in supporting ROS production through its coupling with NOX2. Notably, following cranial window surgery and inoculation with mCherry-GL261 tumor cells, we observed that the windows of *Cx3cr1^gfp/+^*mice often appeared blurrier with excessive macrophage accumulation, while the windows of *Hvcn1^-/-^*:*Cx3cr1^gfp/+^* mice tended to remain clearer. This blurriness could be attributed to significant inflammation, though it remains unclear whether the inflammation is a response to the tumor or its exact nature. Notably, the observed inflammation is unlikely to be due to surgical infection, as all surgeries were performed under strict sterile conditions, and infections would typically result in the loss of the cranial window cap, preventing further imaging. Since this phenotype is associated with accelerated tumor progression, future studies will be needed to identify the specific triggers behind this inflammatory response.

Another important finding in our study is that Hv1 deficiency significantly enhances the antigen-presenting abilities of infiltrating macrophages, ultimately boosting both CD4^+^ and CD8^+^ T-cell tumoricidal responses. This increase in antigen presentation may be due to enhanced uptake of exogenous tumor antigens in macrophages, as we observed upregulation of endocytosis and phagocytosis-related markers, including *P2y6* ^41^ and *Trem2* ^31, 42, 43^, when Hv1 is depleted. Interestingly, previous studies showed that Trem2 depletion impairs the uptake of tumor debris ^31, 43^ and subsequently weakens MHC class II-associated CD4^+^ anti-tumor responses ^31^. Our current findings complement this work by showing that *Trem2* expression increases in the absence of Hv1, although further studies are needed to investigate how Hv1 regulates TREM2 expression.

The enhanced antigen presentation may also be attributed to Hv1’s role in regulating cellular pH, which influences various cellular processes, particularly the acidification of lysosomal and endosomal compartments ^27^. However, the relationship between phagosomal pH and antigen presentation remains complex and controversial, differing across various myeloid cell types and disease contexts ^44, 45, 46^. Hv1 has also been implicated in sustaining calcium entry ^47^, which could further influence monocyte/macrophage polarization through downstream signaling pathways.

It is important to recognize that while Hv1 is not the only proton regulator, its distinctive expression in immune cells suggests that it may not be as broadly targeted as other ion channels in future clinical applications. Other channels, such as the sodium-hydrogen exchanger 1 (NHE1) ^48, 49^ and the vacuolar-type H^+^ ATPase (V-ATPase) ^50^, also play pivotal roles in managing proton dynamics in tumor-associated macrophages. Depleting Hv1 could impact the functioning of these channels, necessitating further research to understand how they might compensate for proton regulation. Moreover, the identification of Hv1 antagonists, such as the C6 peptide which specifically inhibits human voltage-gated proton channels (hHv1) ^51^, marks a significant advancement. With these tools now available, our research provides a foundation for potentially transformative approaches in targeting Hv1 in oncological contexts, particularly in modulating tumor immunity.

## MATERIALS AND METHODS

### Animals

The *Hvcn1^-/-^* mouse line was originally generated by Dr. David E. Clapham lab at the Harvard Medical School and was maintained by Dr. Long-Jun Wu laboratory in Mayo Clinic and University of Texas Health Science Center at Houston. *Hv1-HA-mCherry* and *Hvcn1^flox/flox^* mouse line were generated by Biocytogen (Beijing, China) and bred at Dr. Long-Jun Wu’s laboratory. Wildtype (#000664), *Cx3cr1^creER^* (#021160), *Cx3cr1^gfp^* (#005582), *Ccr2^creER^* (#035229) mice were purchased from Jackson Laboratory and then bred at institutions. All animals were housed under standard conditions (21 - 22 °C; 55% humidity) in individually ventilated cages, with a 12-h light/dark cycle and *ad libitum* access to food and water. Male and female mice aged between 8 to 14 weeks were used in the studies. All experimental procedures were approved by the Mayo Clinic’s and UT Health Houston’s Institutional Animal Care and Use Committee (IACUC).

### Tumor cell culture

Murine GL261 glioma inoculation of C57BL/6 mice is a well-established experimental model of human glioblastoma ^52^. GL261 is a syngeneic mouse model of glioblastoma in C57BL/6 mice that does not require an immunodeficient host (Jacobs, Valdes et al. 2011). The GL261 cell line transduced with firefly luciferase (GL261-luc) for *in vivo* monitoring of tumor kinetics was kindly provided by the laboratory of Dr. Aaron J. Johnson (Mayo Clinic, Rochester, MN). The GL261-luc cell line transduced with mCherry (GL261-luc-mCherry) for *in vivo* two-photon imaging was kindly provided by the laboratory of Dr. Alfredo Quiñones-Hinojosa (Mayo Clinic, Jacksonville, FL). Cells were grown in Dulbecco’s modified Eagle medium (DMEM) (Gibco, #11965092) with 10% fetal bovine serum (FBS) (Sigma-Aldrich, #F2442) and 1% Penicillin-Streptomycin (Gibco, #15140122), in a 37 °C humidified incubator with 5% CO_2_. For tumor inoculation, cells were dissociated with TrypLE™ Express (Gibco, # 12605010) and resuspended in phosphate buffered saline (PBS) to achieve the desired concentration for research purposes.

### Inoculation of GL261

Under isoflurane anesthesia, a 0.5-cm longitudinal incision was made on the scalp, and a burr hole was drilled using a high-speed dental drill (ML: ±1.5; AP: ^+^1.5). Using a stereotactic frame, the needle of a Hamilton syringe was then lowered 3.5 mm into the striatum and a total of 5 × 10^4^ GL261-luc cells in 1-2 μL were injected, as previously described ^53^. Similarly, for 73C inoculation, a total of 1 × 10^4^ cells in 1-2 μL were injected. The wound was closed using 6-0 ETHILON® Nylon Suture (Ethicon, #1660G).

To monitor the tumor burden in GL261-luc-bearing mice, bioluminescence imaging was used as previously described ^53^. Mice were intraperitoneally (IP) injected with 200 μL of 15 mg/μL D-Luciferin in PBS (Goldbio, #LUCK-1G), and anaesthetized with 2% isoflurane during imaging. Mice were scanned using the IVIS Spectrum system (Xenogen Corp.) at Mayo Clinic, running Living Image software. To evaluate the tumor burden, the mouse brains were dissected, tumor hemispheres were weighted.

For survival study, the mice were closely monitored, and when they reached a humane endpoint, showing signs such as hunching, bulging head, or weight loss of approximately 20%, they were euthanized. Additionally, the survival of WT mice with 50,000 GL261 tumor cell inoculation consistently remains around 4 weeks, which has been set as a reliable control for experiments.

### *In vivo* two-photon imaging

Craniotomy and tumor inoculation were performed previously described ^54^ ^31, 55^. In brief, under isoflurane anesthesia (3% induction, 1.5-2% maintenance), a circular craniotomy (<5 mm diameter) was made over somatosensory cortex with the center at about -2.5 posterior and +2 lateral to bregma. A total of 1-2 × 10^3^ GL261-luc-mCherry cells in 0.3 μL were injected into cortex. A circular glass coverslip (4 mm diameter, Warner) was secured over the craniotomy using dental cement (Tetric EvoFlow). A four-point headbar (NeuroTar) was secured over the window using dental cement.

### Generation and culture of bone marrow-derived macrophages (BMDMs)

L-929 fibroblasts (Sigma-Aldrich, 85011425-1VL) were cultured in GlutaMaxTM DMEM (Gibco, 10569010), supplemented with 10% FBS and 1% Penicillin-Streptomycin, to serve as a source of macrophage colony-stimulating factors (M-CSF). Once a sufficient quantity of L-929 culture supernatant was obtained, bone marrow was harvested from WT and *Hvcn1^-/-^* mice’s femurs and tibias, and cultured in GlutaMaxTM DMEM containing 20% L-929 fibroblast cell culture supernatant which supplied the necessary M-CSF, 10% FBS and 1% Penicillin-Streptomycin. The freshly obtained bone marrow cells underwent a 7-day differentiation period to develop into BMDMs, with the culture medium refreshed every 72 hours.

### *In vitro* antigen presentation assay

BMDMs were reseeded into 48-well plates at a density of 0.25 × 10^6^ cells per well. The cells were then exposed to ovalbumin (OVA) (10 ug/ml; sigma A5503) overnight, followed by treatment with 100 ng/ml LPS for overnight. Subsequently, the BMDMs were washed with fresh medium and co-cultured for a period of 48 hours with purified CellTrace™ Far Red (C34572, Invitrogen)-labeled naïve OT-I^+^ CD8^+^ T cells at a 1:1 ratio. T cells were subjected to flow cytometry analysis for flow cytometry, with a focus on assessing dye dilution in viable CD3^+^ CD8^+^ lymphocytes.

### CD11b isolation and bulk RNA sequencing

After perfusing with cold PBS, the tumor was carefully dissected from the affected hemisphere and processed using the Adult Brain Dissociation Kit (130-107-677, Miltenyi Biotec) for digestion, resulting in a single-cell suspension. We then employed the EasySep™ Mouse CD11b Positive Selection Kit II (#18970, STEMCELL) for the targeted isolation of infiltrating myeloid cells. Post-isolation, these cells were concentrated by centrifugation at 1,800 g for 6 minutes at 4 °C. Subsequently, the cell pellet was lysed, and RNA extraction was performed using the RNeasy Micro Kit (#74004, QIAGEN). The RNA obtained was forwarded to BGI for comprehensive quality assessment, sequencing, and pipeline analysis.

### Spectrum flow cytometry

Mice were perfused with 40 mL of 1 × PBS through intracardiac administration. After perfusion, tumor hemispheres were processed using a previously published protocol for enriching brain infiltrating immune cells, using the Dounce Homogenizer followed by centrifugation on a 30% Percoll (Sigma, P1644-1L) gradient ^56^. After staining and fixation, samples were assessed by a spectral flow cytometer (Cytek Aurora, Cytek Biosciences) equipped with SpectroFlo software (Cytek Biosciences). Acquired flow cytometry results were analyzed by FlowJo software (BD Life Sciences).

### Spatial RNA sequencing (10X Visium)

In brief, after intracardially perfused with 40 ml cold PBS, tumor hemispheres were dissected and post-fixed in 4% PFA overnight at 4 ° C. Samples were then processed by the following steps using the automatic tissue processing machine: 70% ethanol (1 hr), 80% ethanol (1 hr), 95% ethanol (30 min), 95% ethanol (30 min), 95% ethanol (30 min), 100% ethanol (30 min), 100% ethanol (30 min), 100% ethanol (45 min), Xylene (60 min), Xylene (60 min), Paraffin (45 min, 60° C), Paraffin (45 min, 60° C), Paraffin (60 min, 60° C), Paraffin (60 min, 60° C). The tissues were then embedded in paraffin to create a paraffin block. The paraffin blocks were then cut using a microtome to generate thin sections of tissue for RNA quality assessment and 10X Visium preparation. After RNA quality assessment (DV200 > 50%), the FFPE tissue block was then sectioned by a microtome at 6 µm to generate appropriately sized sections for Visium slides. The slides dried out for 3 h at 42°C (10X Genomics, CG000408). When slices were completely dried, slides proceeded to deparaffinization, decrosslinking and immunofluorescence staining protocols according to the manufacturer’s protocol with recommended reagents (10X Genomics, 1000339 and 1000251; CG000410). After tissue imaging, the slides were proceeded immediately to Visium Spatial Gene Expression based on User Guide (10X Genomics, CG000407), including probe hybridization, probe ligation, probe release & extension. Library generation commenced with the amplification of eluted probes using real-time qPCR to ascertain the optimal number of amplification cycles. The qPCR protocol commenced with an initial denaturation step lasting 3 minutes at 98°C, succeeded by 25 cycles, each comprising a 5-second duration at 98°C and a 30-second duration at 63°C. The requisite number of PCR cycles for library amplification was established based on the qPCR outcomes, employing 16-19 cycles for the amplification of the entire set of eluted probes, utilizing a 10X dual index kit. Subsequently, the amplified libraries were purified with SPRI select beads (Beckman Coulter). Quantification of the final library was conducted using a TapeStation 4200 D1000 Screen Tape (Agilent) and Qubit (Invitrogen). Sequencing was performed employing paired-end reads of 101 bp, utilizing either the NextSeq2000 P2 or NovaSeq6000 SP at Mayo Clinic Genome Analysis Core (Rochester, MN).

---

**Supplementary figure 1.**
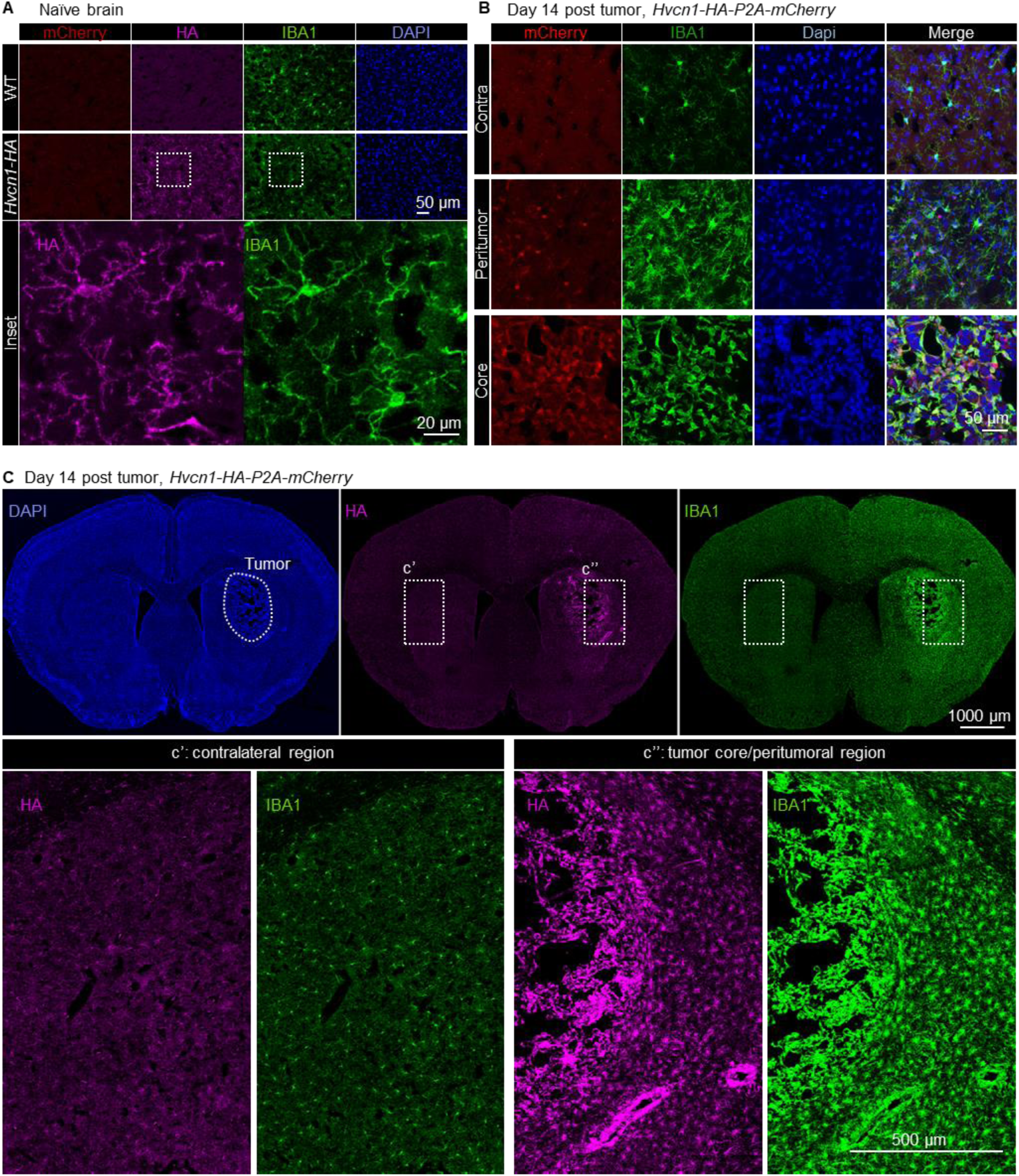
Mapping Hv1 expression in naïve and tumor brains using a novel transgenic mouse line. A. Representative immunostaining displaying of endogenous mCherry, HA, IBA1, and DAPI in the naïve brain. WT served as a negative control for mCherry and HA staining (scale bar: 50 µm). An inset demonstrates co-localization of HA with IBA1^+^ cells (scale bar: 20 µm). B. Utilizes the same staining protocol as in (A) for brains 14 days after tumor cell inoculation (scale bar: 50 µm). C. Displays the overall staining pattern 14 days after tumor inoculation (scale bar: 1000 µm). Under the same confocal setting, inset C’ (left) displays ramified microglia with relatively uniform and dim HA staining in the contralateral region. In contrast, inset C’’ (right) illustrates activated microglia in response to glioma, where infiltrating myeloid cells exhibit more intense Hv1 staining (scale bar: 500 µm).

**Supplementary Figure 2.**
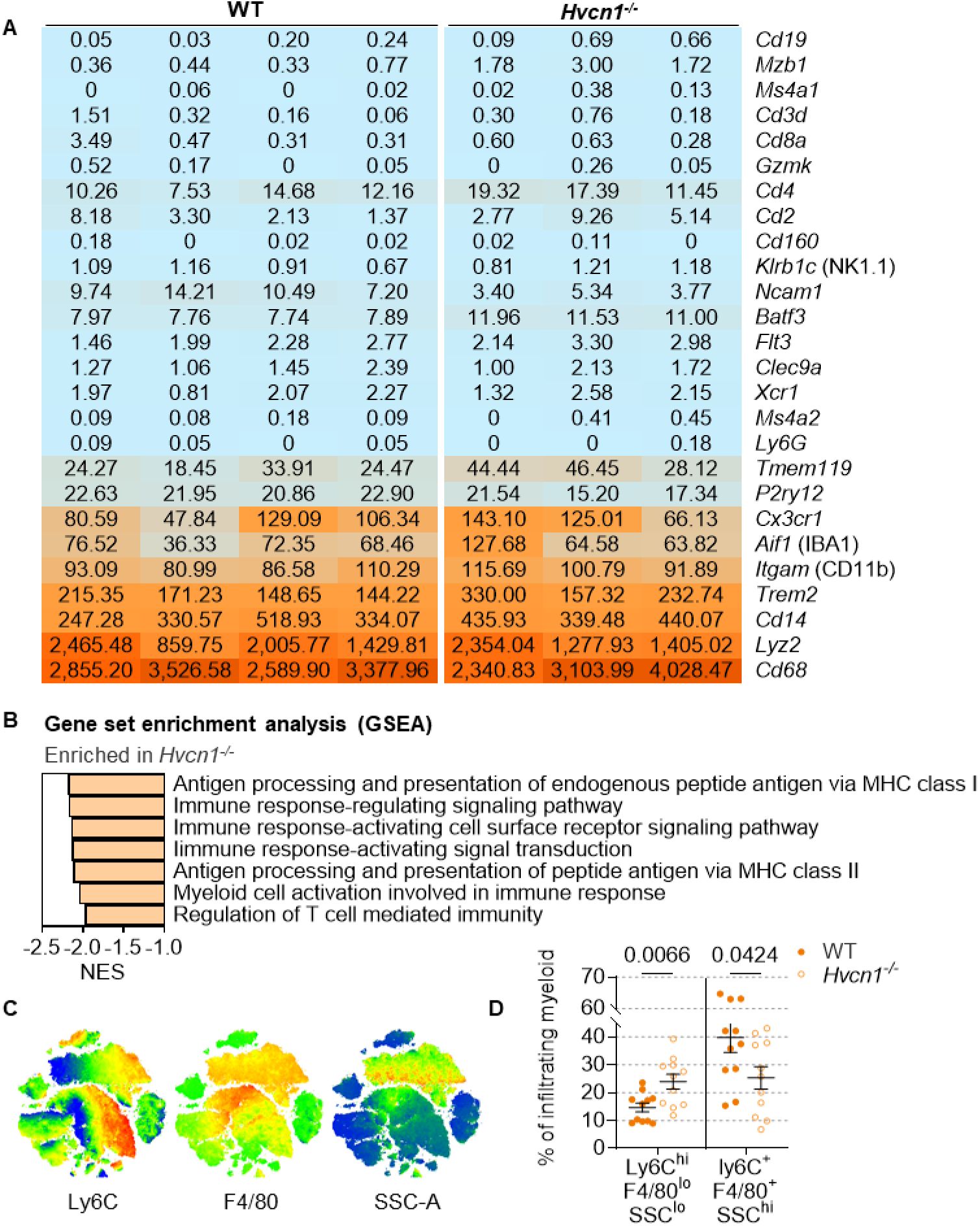
Characterization of infiltrating myeloid cells through both transcriptomic profiling and protein marker assessment. A. Selected marker genes confirm the predominance of infiltrating myeloid cells in the sorted populations. B. GSEA reveals strong myeloid activation in *Hvcn1^-/-^*, particularly mechanisms involved in antigen presentation and immune signaling regulation. C. Displays Ly6C, F4/80, and SSC-A on a tSNE map, illustrating the distribution and clustering of myeloid cells. D. Analysis of the percentages of different myeloid populations in wild-type (WT) versus *Hvcn1^-/-^*. Data were assessed for normal distribution using Shapiro-Wilk test are presented as mean ± SEM for normally distributed data or medium ± interquartile range for non-normally distributed data. P-values were determined using two-tailed Student t-tests for normally distributed data, and Mann-Whitney test for non-normally distributed data.

**Supplementary Figure 3.**
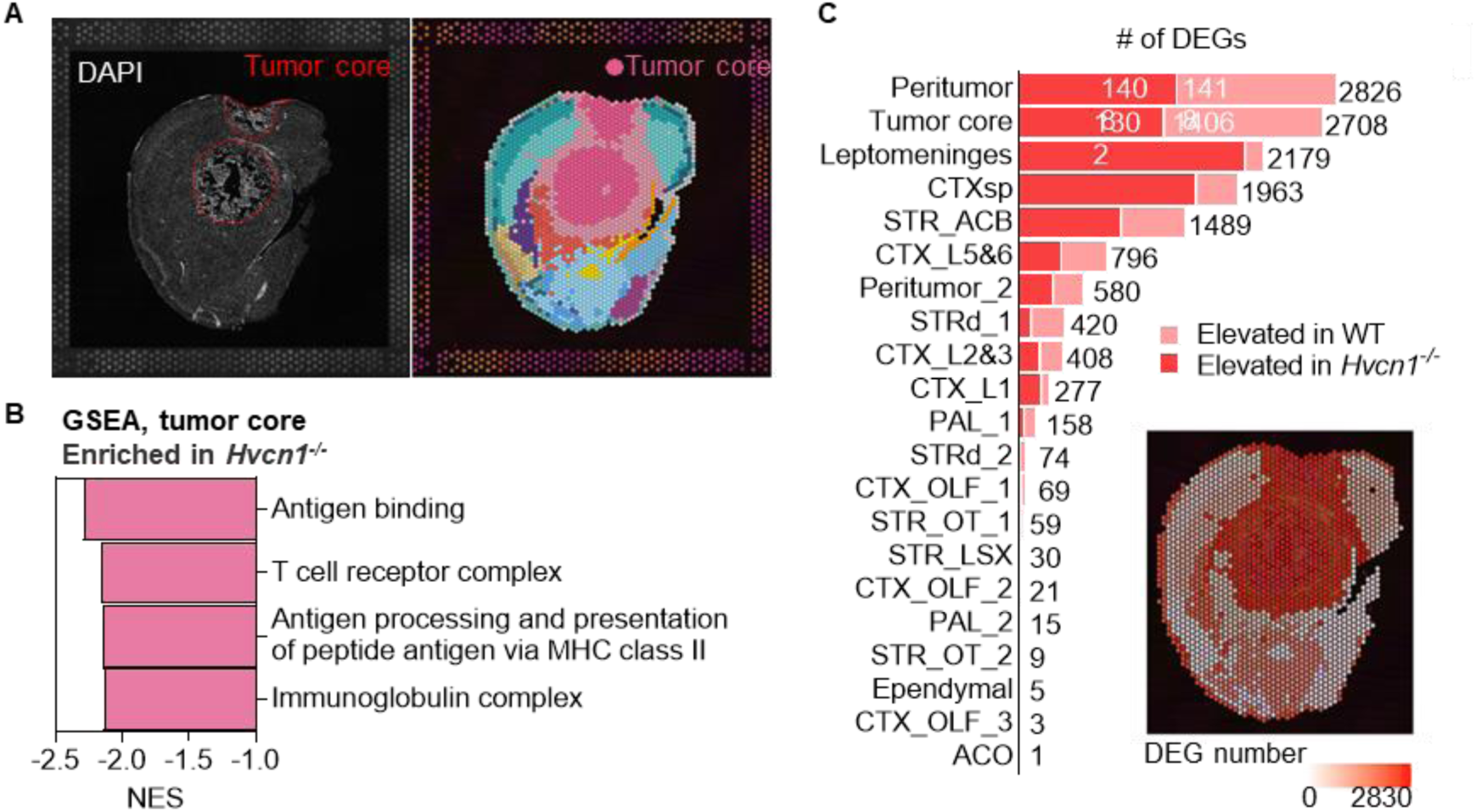
Spatial Transcriptomics Analysis. A. Representative images show spatial RNA sequencing closely aligned with the dense DAPI staining that marks the tumor region. B. GSEA results for the tumor core highlight several immune-related pathways enriched in *Hvcn1^-/-^*. C. Number of DEGs across multiple annotated regions are presented.

## REFERENCE

1. Zheng, J., Murugan, M., Wang, L. & Wu, L.-J. Microglial voltage-gated proton channel Hv1 in spinal cord injury. Neural Regeneration Research 17 (2022).

2. Koch, H.P. et al. Multimeric nature of voltage-gated proton channels. Proceedings of the National Academy of Sciences of the United States of America 105, 9111–9116 (2008).

3. Tombola, F., Ulbrich, M.H. & Isacoff, E.Y. The voltage-gated proton channel Hv1 has two pores, each controlled by one voltage sensor. Neuron 58, 546–556 (2008).

4. Lee, S.Y., Letts, J.A. & Mackinnon, R. Dimeric subunit stoichiometry of the human voltage-dependent proton channel Hv1. Proceedings of the National Academy of Sciences of the United States of America 105, 7692–7695 (2008).

5. Ramsey, I.S., Moran, M.M., Chong, J.A. & Clapham, D.E. A voltage-gated proton-selective channel lacking the pore domain. Nature 440, 1213–1216 (2006).

6. Sasaki, M., Takagi, M. & Okamura, Y. A voltage sensor-domain protein is a voltage-gated proton channel. Science 312, 589–592 (2006).

7. Wu, L.J. Voltage-gated proton channel HV1 in microglia. Neuroscientist 20, 599–609 (2014).

8. DeCoursey, T.E. The Voltage-Gated Proton Channel: A Riddle, Wrapped in a Mystery, inside an Enigma. Biochemistry 54, 3250–3268 (2015).

9. Lishko, P.V., Botchkina, I.L., Fedorenko, A. & Kirichok, Y. Acid extrusion from human spermatozoa is mediated by flagellar voltage-gated proton channel. Cell 140, 327–337 (2010).

10. Hondares, E. et al. Enhanced activation of an amino-terminally truncated isoform of the voltage-gated proton channel HVCN1 enriched in malignant B cells. Proc Natl Acad Sci U S A 111, 18078–18083 (2014).

11. Wang, Y. et al. Specific expression of the human voltage-gated proton channel Hv1 in highly metastatic breast cancer cells, promotes tumor progression and metastasis. Biochem Biophys Res Commun 412, 353–359 (2011).

12. Wang, Y., Li, S.J., Wu, X., Che, Y. & Li, Q. Clinicopathological and biological significance of human voltage-gated proton channel Hv1 protein overexpression in breast cancer. J Biol Chem 287, 13877–13888 (2012).

13. Wu, L.J. et al. The voltage-gated proton channel Hv1 enhances brain damage from ischemic stroke. Nat Neurosci 15, 565–573 (2012).

14. Murugan M., et al. Voltage-gated proton channel Hv1 promotes secondary damage following spinal cord injury Molecular Brain (2020).

15. Li, X. et al. Microglial Hv1 exacerbates secondary damage after spinal cord injury in mice. Biochem Biophys Res Commun (2020).

16. Li, X. et al. Deficiency of the microglial Hv1 proton channel attenuates neuronal pyroptosis and inhibits inflammatory reaction after spinal cord injury. J Neuroinflammation 17, 263 (2020).

17. Li, Y., et al. The voltage-gated proton channel Hv1 plays a detrimental role in contusion spinal cord injury via extracellular acidosis-mediated neuroinflammation Brain Behav and Immun (2020).

18. Liu, J. et al. Microglial Hv1 proton channel promotes cuprizone-induced demyelination through oxidative damage. J Neurochem 135, 347–356 (2015).

19. Ritzel, R.M. et al. Proton extrusion during oxidative burst in microglia exacerbates pathological acidosis following traumatic brain injury. Glia 69, 746–764 (2021).

20. Peng, J. et al. The voltage-gated proton channel Hv1 promotes microglia-astrocyte communication and neuropathic pain after peripheral nerve injury. Mol Brain 14, 99 (2021).

21. Pombo Antunes, A.R., et al. Single-cell profiling of myeloid cells in glioblastoma across species and disease stage reveals macrophage competition and specialization. Nat Neurosci 24, 595–610 (2021).

22. Kilian, M. et al. MHC class II-restricted antigen presentation is required to prevent dysfunction of cytotoxic T cells by blood-borne myeloids in brain tumors. Cancer Cell (2022).

23. Chen, D. et al. CTLA-4 blockade induces a microglia-Th1 cell partnership that stimulates microglia phagocytosis and anti-tumor function in glioblastoma. Immunity 56, 2086–2104 e2088 (2023).

24. Veglia, F., Sanseviero, E. & Gabrilovich, D.I. Myeloid-derived suppressor cells in the era of increasing myeloid cell diversity. Nat Rev Immunol 21, 485–498 (2021).

25. Gill, B.J. et al. MRI-localized biopsies reveal subtype-specific differences in molecular and cellular composition at the margins of glioblastoma. Proc Natl Acad Sci U S A 111, 12550–12555 (2014).

26. Klemm, F. et al. Interrogation of the Microenvironmental Landscape in Brain Tumors Reveals Disease-Specific Alterations of Immune Cells. Cell 181, 1643–1660 e1617 (2020).

27. Montes-Cobos, E. et al. Voltage-Gated Proton Channel Hv1 Controls TLR9 Activation in Plasmacytoid Dendritic Cells. J Immunol 205, 3001–3010 (2020).

28. Peng, J. et al. Microglia and monocytes synergistically promote the transition from acute to chronic pain after nerve injury. Nat Commun 7, 12029 (2016).

29. Parkhurst, C.N. et al. Microglia promote learning-dependent synapse formation through brain-derived neurotrophic factor. Cell 155, 1596–1609 (2013).

30. Flores-Toro, J.A. et al. CCR2 inhibition reduces tumor myeloid cells and unmasks a checkpoint inhibitor effect to slow progression of resistant murine gliomas. Proc Natl Acad Sci U S A 117, 1129–1138 (2020).

31. Zheng, J. et al. TREM2 mediates MHCII-associated CD4+ T-cell response against gliomas. Neuro Oncol 26, 811–825 (2024).

32. Darmanis, S. et al. Single-Cell RNA-Seq Analysis of Infiltrating Neoplastic Cells at the Migrating Front of Human Glioblastoma. Cell reports 21, 1399–1410 (2017).

33. Chen, Z. et al. Cellular and Molecular Identity of Tumor-Associated Macrophages in Glioblastoma. Cancer Res 77, 2266–2278 (2017).

34. Ochocka, N. et al. Single-cell RNA sequencing reveals functional heterogeneity of glioma-associated brain macrophages. Nat Commun 12, 1151 (2021).

35. Lein, E.S. et al. Genome-wide atlas of gene expression in the adult mouse brain. Nature 445, 168–176 (2007).

36. Kruse, B. et al. CD4(+) T cell-induced inflammatory cell death controls immune-evasive tumours. Nature 618, 1033–1040 (2023).

37. Chen, D. et al. CTLA-4 blockade induces CD4(+) T cell IFNgamma-driven microglial phagocytosis and anti-tumor function in glioblastoma. Immunity (2023).

38. Li, Y. et al. The voltage-gated proton channel Hv1 plays a detrimental role in contusion spinal cord injury via extracellular acidosis-mediated neuroinflammation. Brain Behav Immun 91, 267–283 (2021).

39. DeNardo, D.G. & Ruffell, B. Macrophages as regulators of tumour immunity and immunotherapy. Nat Rev Immunol 19, 369–382 (2019).

40. Groth, C. et al. Immunosuppression mediated by myeloid-derived suppressor cells (MDSCs) during tumour progression. Br J Cancer 120, 16–25 (2019).

41. Umpierre, A.D. et al. Microglial P2Y(6) calcium signaling promotes phagocytosis and shapes neuroimmune responses in epileptogenesis. Neuron 112, 1959–1977 e1910 (2024).

42. Bosco, D.B. et al. Microglial TREM2 promotes phagocytic clearance of damaged neurons after status epilepticus. Brain Behav Immun 123, 540–555 (2024).

43. Peshoff, M.M. et al. Triggering receptor expressed on myeloid cells 2 (TREM2) regulates phagocytosis in glioblastoma. bioRxiv, 2023.2004.2005.535792 (2023).

44. Bagaitkar, J. et al. NADPH oxidase activation regulates apoptotic neutrophil clearance by murine macrophages. Blood 131, 2367–2378 (2018).

45. Alloatti, A., Kotsias, F., Magalhaes, J.G. & Amigorena, S. Dendritic cell maturation and cross-presentation: timing matters! Immunol Rev 272, 97–108 (2016).

46. Mantegazza, A.R. et al. NADPH oxidase controls phagosomal pH and antigen cross-presentation in human dendritic cells. Blood 112, 4712–4722 (2008).

47. El Chemaly, A., et al. VSOP/Hv1 proton channels sustain calcium entry, neutrophil migration, and superoxide production by limiting cell depolarization and acidification. J Exp Med 207, 129–139 (2010).

48. Hasan, M.N. et al. Blocking NHE1 stimulates glioma tumor immunity by restoring OXPHOS function of myeloid cells. Theranostics 11, 1295–1309 (2021).

49. Guan, X. et al. Blockade of Na/H exchanger stimulates glioma tumor immunogenicity and enhances combinatorial TMZ and anti-PD-1 therapy. Cell Death Dis 9, 1010 (2018).

50. Liu, N. et al. Lactate inhibits ATP6V0d2 expression in tumor-associated macrophages to promote HIF-2alpha-mediated tumor progression. J Clin Invest 129, 631–646 (2019).

51. Zhao, R., Shen, R., Dai, H., Perozo, E. & Goldstein, S.A.N. Molecular determinants of inhibition of the human proton channel hHv1 by the designer peptide C6 and a bivalent derivative. Proc Natl Acad Sci U S A 119, e2120750119 (2022).

52. Haddad, A.F. et al. Mouse models of glioblastoma for the evaluation of novel therapeutic strategies. Neurooncol Adv 3, vdab100 (2021).

53. Ayasoufi, K. et al. Brain cancer induces systemic immunosuppression through release of non-steroid soluble mediators. Brain 143, 3629–3652 (2020).

54. Liu, Y.U. et al. Neuronal network activity controls microglial process surveillance in awake mice via norepinephrine signaling. Nature neuroscience 22, 1771–1781 (2019).

55. Eyo, U.B. et al. P2Y12R-Dependent Translocation Mechanisms Gate the Changing Microglial Landscape. Cell reports 23, 959–966 (2018).

56. Cumba Garcia, L.M., Huseby Kelcher, A.M., Malo, C.S. & Johnson, A.J. Superior isolation of antigen-specific brain infiltrating T cells using manual homogenization technique. J Immunol Methods 439, 23–28 (2016).

